# Ion selectivity as a biophysical determinant of action-potential dynamics

**DOI:** 10.64898/2026.11.13.766596

**Authors:** Ada Y. Chen, Bernard R. Brooks, Ana Damjanovic

## Abstract

Ion selectivity is a fundamental property of voltage-gated ion channels, yet its contribution to action-potential dynamics has received less attention than channel conductance and gating kinetics. Here, we investigate how Na^+^/K^+^ permeability ratios influence membrane excitability, extending a multiscale framework that connects molecular properties of ion channels to neuronal electrical activity. We employ a simplified membrane depolarization model and a modified Hodgkin-Huxley model in which channel reversal potentials are determined by the Goldman-Hodgkin-Katz equation, allowing selectivity to be varied independently of conductance and gating. We find that decreasing Na_v_ selectivity reduces the inward driving force, slowing the action-potential upstroke and lowering its peak amplitude. This dependence is nonlinear: action-potential amplitude is more sensitive to changes at lower Na_v_ selectivity and approaches a plateau as selectivity increases. Reduced K_v_ selectivity has qualitatively different consequences, shifting the equilibrium membrane potential toward more positive values and producing distinct dynamical regimes, including spontaneous firing and sustained depolarization with suppressed firing. Changes in Na_v_ conductance affect action-potential dynamics through a distinct mechanism. Together, these results identify ion selectivity as an independent biophysical determinant of neuronal excitability and establish a quantitative connection between pore-level permeability and cellular electrical behavior. This approach provides a basis for incorporating molecular changes in ion selectivity into multiscale models of neuronal signaling.

## Introduction

Selectivity is a fundamental property of biological ion channels. Ion channels span a broad range of selectivities, from channels selective for one ion species, such as voltage-gated sodium (Na_v_) and potassium (K_v_) channels, to channels that conduct several ion species (such as many synaptic channels selective for cations), to anion selective channels, and relatively non-selective channels. The selectivity of a channel for one ion relative to another can be quantified by their relative permeabilities, for example by the permeability ratio *P*_Na_/*P*_K_. Voltage gated Na_v_ and K_v_ are central for action potential generation. When the membrane depolarizes, Na_v_ channels open rapidly and allow Na^+^ ions to flow down their electrochemical gradient, producing an inward current. This inward current further depolarizes the membrane and opens additional sodium channels, creating a positive feedback loop that drives the rapid rise of the action potential (1, 2). Voltage-gated K_v_ channels activate more slowly in response to membrane depolarization and allow K^+^ ions to flow outward, producing an outward current that repolarizes the membrane and helps restore the resting membrane potential. Because of the selectivity of Na_v_ and K_v_ channels, the ionic currents underlying membrane depolarization and repolarization during action potentials are carried predominantly by Na^+^ and K^+^ ions, respectively.

Action potentials exhibit substantial variation in their shape across different cells and organisms, including differences in rise time, peak amplitude, and duration, as well as distinctive features such as kinks and plateaus. These differences are commonly attributed to variations in channel density, channel types, gating kinetics, or ionic concentration gradients (2–4). Differences in ion selectivity could provide an additional biophysical factor influencing action-potential waveform, but the effects of varying selectivity independently of other channel properties have received considerably less attention.

The current carried by an ion channel depends on channel conductance, gating, and the electrochemical forces that drive ions across the membrane, i.e., the concentration difference across the membrane and the electrical potential across the membrane. The reversal potential is the membrane voltage at which the net current through the channel is zero. For a channel that is perfectly selective for a single ion species, the reversal potential is given by the Nernst equation. When multiple ion species can permeate the same channel, the reversal potential depends on their concentration gradients and relative ion permeabilities, as described by the Goldman-Hodgkin-Katz (GHK) equation (5, 6). Thus, changes in ion selectivity can shift the channel reversal potential and alter the current flowing through the channel, even when conductance and gating are unchanged.

Ion selectivity in voltage-gated sodium channels arises from specific structural features of the pore. In eukaryotic Na_v_ channels the highly conserved DEKA motif, four different residues: Asp, Glu, Lys, and Ala, line the narrow region of the pore (7). Bacterial Na_v_ channels instead contain a ring of four glutamate residues. Although these structural features are known to be critical for Na^+^ selectivity, the physical mechanism by which bacterial Na_v_ channels discriminate between Na^+^ and K^+^ remains under debate, with ion blockage (8), pKa shifts (9), multi-ion effects (10), steric effects (11), and ion-induced filter rearrangement (12) all implicated. In eukaryotic Na_v_ channels, the lysine residue within the DEKA motif is critical for Na^+^/ K^+^ discrimination (7), while proposed selectivity mechanisms include electrostatic steering by lysine toward acidic residues that preferentially coordinate Na^+^ over K^+^ (13, 14), multi-ion complexation (15), and changes in lysine protonation state (16), although the detailed mechanism remains unresolved and simulations do not always reproduce experimentally observed Na^+^/ K^+^ selectivity (17). In K_v_ channels, the selectivity filter contains a highly conserved sequence whose backbone carbonyl oxygens form a series of closely spaced ion-binding sites that preferentially coordinate K^+^ over Na^+^ (18). Interestingly, ion selectivity is not a fixed property shared uniformly by all Na_v_ or K_v_ channels: experimentally measured permeability ratios vary among channel types and can be substantially altered by mutations within or near the selectivity filter. Reported *P*_Na_/*P*_K_ ratios for eukaryotic Na_v_ channels include 3.5 - 12.8 for squid giant axon channels (19), approximately 12.5 - 33.3 for rat skeletal-muscle Na_v_1.4 (7, 20, 21), and 5.3 - 5.9 for human Na_v_1.7 (22). For bacterial Na^+^ channels, measurements include approximately 1.04 - 171 across experimental preparations for NaChBac (23–25), reported lower bounds ranging from 5 to 25 for Na_v_Ms (26, 27), and 3.7 - 4.5 for pore-only Na_v_Sp1p constructs (28, 29). K^+^ channels can exhibit *P*_K_/*P*_Na_ ratios approaching 1000 (30, 31). Specific K_v_ measurements include 25 for K_v_1.2 (32), approximately 27.0 - 66.7 for K_v_2.1, including human-channel measurements (33–35), 71.4 for human K_v_3.1 (35), and approximately 11 to >400 for Drosophila Shaker, depending on inactivation state (36, 37). These estimates depend on experimental conditions, channel state, and preparation.

The observed variability in Na^+^/K^+^ permeability ratios raises questions about what range of permeability ratios is compatible with normal electrical function and whether different degrees of selectivity have distinct consequences for membrane excitability. Despite extensive work on the molecular and structural determinants of ion selectivity, the consequences of Na^+^/K^+^ selectivity for electrical signaling remain less well understood. In particular, it is not clear how variations in Na^+^/K^++^ permeability ratios influence the dynamics of the action potential, including its rise time and peak amplitude. Recent studies have begun to explore multiscale connections between molecular simulations of ion channels and models of membrane excitability (38), highlighting the importance of linking pore-level properties to cellular electrical behavior. However, the specific role of Na^+^/K^+^ selectivity in shaping action potential dynamics has not been systematically investigated. Such an approach would provide a multiscale framework for linking molecular changes in ion-channel selectivity to their effects on neuronal excitability and action-potential dynamics.

Here we build on our previous molecular studies of sodium channel selectivity (9) by examining how Na^+^/K^+^ permeability ratios influence electrical signaling. By independently varying Na^+^/K^+^ permeability ratios in Na_v_ and K_v_ channels, and Na_v_ channel conductance, we determine how these biophysical parameters influence membrane depolarization, repolarization, and action-potential dynamics. This framework allows us to isolate the contribution of ion selectivity from other channel properties and to connect pore-level permeability to cellular electrical behavior.

## Methods

### Model overview

Two computational models were used to investigate how Na^+^-to-K^+^ selectivity influences depolarization and action-potential generation. The first model, referred to below as the depolarization-only model, was designed to isolate the effect of mixed Na^+^ and K^+^ permeation through Na_v_ channels on the early rise of membrane voltage. The second model was a modified Hodgkin-Huxley model, in which classical Hodgkin-Huxley gating kinetics were retained, whereas channel reversal potentials were reformulated by Goldman-Hodgkin-Katz (GHK) equation to depend on Na^+^ and K^+^ permeability ratios and ionic concentrations. In both models, selectivity was controlled through the ratio P_Na_/P_K_.

### Depolarization-only model

The depolarization-only model describes a local membrane region associated with mixed-permeability conduction pathway through Na_v_ channels. The model tracks membrane voltage in discrete time steps. Selectivity was introduced through a mixed Na^+^/K^+^ reversal potential calculated with the GHK expression:

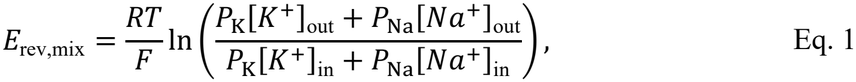

where P_Na_ and P_K_ denote the Na^+^ and K^+^ permeabilities of the Na_v_ channel and, in practice, are implemented as relative permeability weights. R is the gas constant, T is absolute temperature, and F is Faraday’s constant.

The driving voltage difference of the mixed current was then written as

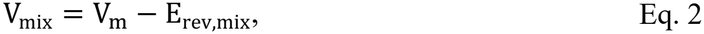

where V_m_ is membrane voltage. Using this driving force, and assuming an Ohmic current - voltage relationship with a constant conductance, the total transferred charge during one time step was calculated as

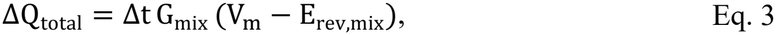

where Δt is the time step and G_mix_ is the effective conductance of the mixed-permeability pathway. In the implementation, Δt was 1 ns.

The membrane voltage update was obtained by converting transferred charge to a voltage change, by modeling the membrane as a plane capacitor, through

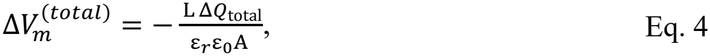

where L is membrane thickness, A is the effective membrane-associated area, ε_0_ is the vacuum permittivity, and ε_r_ is the membrane relative permittivity. We set ε_r_ = 3.2 based on the effective dielectric constant measured for supported dipalmitoylphosphatidylcholine (DPPC) bilayers in aqueous solution (39). This value was adopted as a representative effective permittivity for the lipid bilayer.

The membrane voltage at the next time step was then updated as

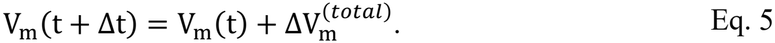

This voltage-update procedure advances the membrane potential iteratively through time. In the Ohmic formulation, however, the total charge transferred through a mixed-permeability pathway cannot be uniquely decomposed into ion-specific Na^+^ and K^+^ contributions. The corresponding changes in Na^+^ and K^+^ concentrations therefore cannot be determined, and all ionic concentrations were held constant throughout the simulation. Consequently, the mixed reversal potential in Eq. 1 was also constant. As a validation of this approximation, we additionally implemented a GHK-current version of the model in which the Na^+^ and K^+^ currents were calculated separately and used to update their intracellular concentrations, while the extracellular concentrations were treated as fixed reservoirs (see Supplementary Methods and Figure S1). The resulting intracellular concentration changes were less than 1%, supporting the constant-concentration approximation used in this Ohmic model (Figure S1D&E).

The conductance of the mixed-permeability pathway G_mix_ was set to 20 pS, a representative single channel conductance value within the approximately 15 - 25 pS range reported for mammalian Na_v_ channels (40–42). This conductance was held fixed across selectivity conditions, while its Na^+^/K^+^ selectivity was represented through the mixed reversal potential. We used this formulation rather than introducing separate Ohmic Na^+^ and K^+^ currents through the same channel because P_Na_/P_K_ does not uniquely specify a pair of constant Na^+^ and K^+^ conductances; according to the GHK current equation (Eq. S1), the relationship between permeability and ionic current also depends on membrane voltage and ion concentrations. This formulation therefore allows Na^+^/K^+^ selectivity to be varied independently while keeping the total channel conductance fixed.

Based on the estimated Na^+^ channel density of approximately 150 channels per µm^2^ in the axon initial segment of cortical layer 5 pyramidal neurons (43), the effective membrane area A associated with each channel was set to 0.007 µm^2^. Membrane thickness was set to a typical value of 5 nm (44). These quantities were used as coarse-grained physical scales rather than as exact anatomical parameters.

Depolarization-only simulations were started from a membrane voltage of −60 mV with extracellular and intracellular Na^+^ concentrations of 145 and 12 mM, respectively, and extracellular and intracellular K^+^ concentrations of 5 and 140 mM, respectively. These concentrations are representative of mammalian neurons, for which typical values are [Na^+^]_out_ = 145 mM, [Na^+^]_in_ = 5 - 15 mM, [K^+^]_out_ = 5 mM, and [K^+^]_in_ = 140 mM (45).

Simulations were run for 10000 steps, corresponding to a total simulation time of 10 µs. The tested selectivity ratios were P_Na_/P_K_ = 100000, 20, 5, and 1. The value 20 was included as a representative physiological selectivity ratio for mammalian Na_v_ channels, for which reported Na^+^/K^+^ permeability ratios generally fall between approximately 13 and 33 (46).

### Classical Hodgkin-Huxley model

The second model was based on the classical Hodgkin-Huxley formalism (1) with membrane capacitance C_m_ equal to 1.0 µF/cm^2^, maximal conductance g_Na_ equal to 120 mS/cm^2^, g_K_ equal to 36 mS/cm^2^, and g_L_ equal to 0.3 mS/cm^2^, and leak reversal potential E_L_ equal to −54.387 mV. The membrane equation was

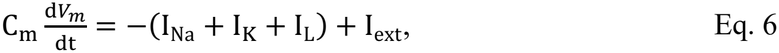

where V_m_ is the membrane voltage, and I_ext_ is the injected current. Eq. 6 was integrated by forward Euler method with a time step of 0.01 ms. Reducing the Hodgkin-Huxley integration time step to 1 ns, matching that of the depolarization-only model, produced no substantive changes in the time courses of the simulated quantities, demonstrating that the chosen time step of 0.01 ms is adequate for the reported results.

The ionic currents were defined as

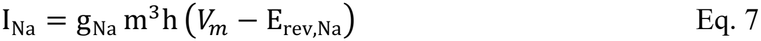

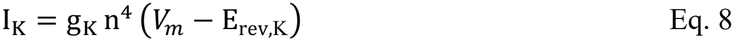

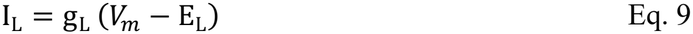

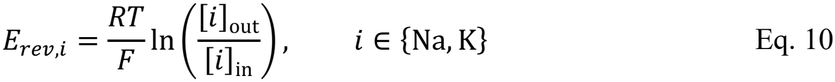

The variables m and h describe Na_v_ activation and availability, respectively, and n describes K_v_ activation; *m*^3^*h* and *n*^4^ are the corresponding gating factors.

These current expressions, Eq. 7 to Eq. 9, use the classical Ohmic approximation: for a fixed open probability, the open-channel current-voltage relationship is assumed to be approximately linear, and the current is proportional to the driving force *V*_*m*_ − *E*_rav_. In the classical Hodgkin-Huxley simulations, *E*_rav,Na_ and *E*_rav,K_ were fixed single-ion reversal potentials calculated from the Nernst equation Eq. 10.

The gating variables obeyed the standard Hodgkin-Huxley equations

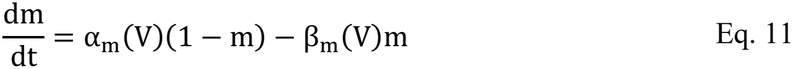

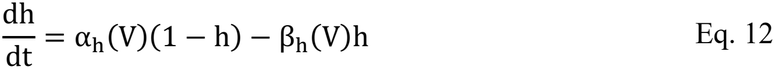

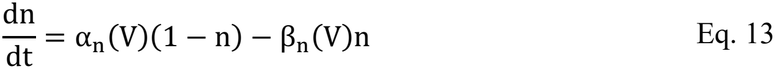

with the voltage-dependent rates

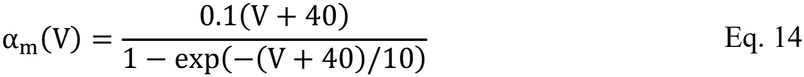

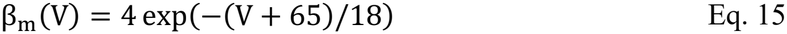

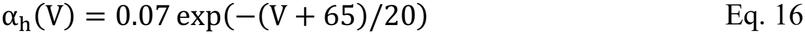

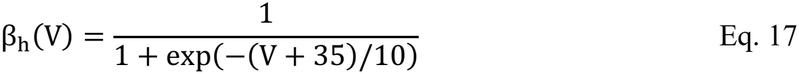

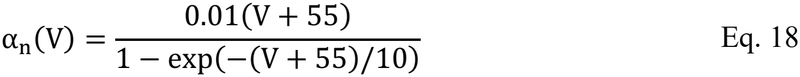

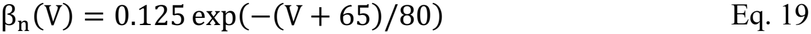

Equations Eq. 11–Eq. 19 were evaluated directly as functions of membrane voltage V_m_. Note that these empirical parameters were originally fitted to the squid giant axon and may not be directly generalizable to other neurons. Here, however, our goal is to isolate the effects of selectivity, conductance, and ionic concentrations (Figure S2), so keeping the gating kinetics fixed is a reasonable simplification.

The classical Hodgkin-Huxley model was parameterized from squid giant axon measurements at 6.3 °C and uses Na^+^ and K^+^ reversal potentials of +50 and −77 mV, respectively, when voltage is expressed in the modern convention with a resting potential of −65 mV (1). To ensure that the mixed GHK reversal potential approached these classical values at the Na^+^- and K^+^-selective limits, effective ion concentrations were selected using the Nernst equation Eq. 10. With [Na^+^]_out_ = 440 mM and [K^+^]_in_ = 400 mM as representative squid axon concentrations (47), the corresponding values [Na^+^]_in_ = 55.17 mM and [K^+^]_out_ = 16.35 mM reproduce E_Na_ = +50 mV and E_K_ = −77 mV at 6.3°C. These should therefore be regarded as model-consistent effective concentrations rather than a single experimentally measured concentration set.

These concentrations differ from those used in the depolarization-only model because the two models serve different purposes. The Hodgkin-Huxley simulations were designed to preserve the reversal potentials and electrophysiological behavior of the classical squid axon model, whereas the depolarization-only model uses representative mammalian ionic conditions and a representative mammalian Na_v_ single-channel conductance to examine the direct effects of Na_v_ channel selectivity. Because comparisons are made across selectivity values within each model, the two models are not required to use identical concentrations or temperatures. Accordingly, comparisons between the two models focus on qualitative trends in the effects of selectivity, rather than quantitative agreement in voltage amplitudes or time scales. Action potentials were elicited under current clamp using a square current pulse of 12 µA/cm^2^ from 10 to 40 ms. In the code, this stimulation protocol was implemented as a time-dependent external current term in Eq. 6.

### Modified Hodgkin-Huxley model for selectivity variations

For the Na^+^ channel selectivity analysis, the Na^+^ channel reversal potential in Eq. 7 was replaced by the mixed Na^+^/K^+^ GHK reversal potential from Eq. 1, whereas the K^+^ channel current in Eq. 8 retained its classical K^+^ reversal potential. Thus, the permeability ratio directly controlled the composition of the current through the Na^+^ channel. The tested Na^+^ channel selectivity values were P_Na_/P_K_ = 100000, 12.8, 5, 1. The value 12.8 was included as a representative physiological Na^+^ channel selectivity ratio based on measurements from squid axon Na^+^ channels under high intracellular K^+^ conditions (19).

For the K^+^ channel selectivity analysis, the K^+^ channel reversal potential in Eq. 8 was replaced by the mixed Na^+^/K^+^ GHK reversal potential from Eq. 1, whereas the Na^+^ channel current in Eq. 7 retained a fixed Na^+^ reversal potential. Thus, the permeability ratio controlled the composition of the current through the K^+^ channel. The tested K^+^ channel selectivity values were P_Na_/P_K_ = 1/100000, 0.03, 0.2, 1. The value 0.03 was included as a representative physiological K^+^ channel selectivity ratio in squid axon (48). Because residual Na^+^ conductance could bias this estimate upward, it may be regarded as an upper limit (49). This value is also consistent with the upper-bound estimate (≲ 0.06) obtained for squid giant axon K^+^ channels (49, 50).

### Hodgkin-Huxley model for Na^+^ channel conductance variation

For the conductance analysis, the classical Ohmic Na^+^ and K^+^ current equations were retained. The maximal Na^+^ channel conductance in Eq. 7 was varied while the Na^+^ channel was kept fully Na^+^-selective. The three conductance values examined were 60, 120, and 180 mS/cm^2^, with 120 mS/cm^2^ being the default. In this way, the effect of changing Na^+^ channel conductance could be examined independently of the Na^+^-to-K^+^ permeability ratio.

### Numerical implementation

All simulations were implemented in Python using NumPy for numerical calculations and Matplotlib for visualization. The depolarization-only model used sequential discrete-time updates of membrane voltage, whereas the classical and modified Hodgkin-Huxley models used explicit forward Euler integration of the membrane and gating equations.

For each Hodgkin-Huxley parameter set, the zero-current equilibrium membrane voltage was calculated by solving *I*_Nav_ + *I*_Kv_ + *I*_*L*_ = 0 with no injected current and the gating variables set to their voltage-dependent steady-state values, *x*_∞_ (*V*) = *α*_*x*_(*V*)/[*α*_*x*_ (*V*) + *β*_*x*_ (*V*)], where *x* ∈ {*m, h, n*}. Roots were obtained using Brent’s bracketed root-finding method. These conditions define a fixed point at which the time derivatives of all four dynamical variables (V, m, h, and n) vanish. Local stability was assessed from the eigenvalues of the full voltage-m-h-n Jacobian, evaluated numerically by central finite differences at each equilibrium voltage. An equilibrium voltage was classified as locally asymptotically stable when all eigenvalues had negative real parts and unstable when at least one had a positive real part. In the linearized equations, perturbation modes evolve as *e*^*λ t*^. A positive real part therefore allows small deviations to grow even though the equilibrium voltage satisfies the current-balance equation. When multiple roots were found, the stable root nearest −65 mV was preferred; if none was stable, the nearest unstable root was retained. To avoid a separate equilibration step, the main simulations were therefore initialized directly at their pre-calculated equilibrium voltages, with the corresponding steady-state gating values. For comparison, we also ran K_v_ simulations from a common initial voltage of −65 mV with a 100 ms pre-stimulus equilibration interval (Figure S4), revealing condition-dependent relaxation toward equilibrium or persistent firing when the equilibrium voltage was unstable.

## Results

### Depolarization-only model

We first used the depolarization-only model to examine how Na^+^-to-K^+^ selectivity affects the early rise of membrane voltage. Figure 1A shows membrane voltage traces for permeability ratios P_Na_/P_K_ of 1, 5, 20 and 100000. Increasing the permeability ratio led to a progressively faster rise of voltage and a larger voltage change over the same timescale. Thus, in this reduced model, stronger Na^+^ selectivity increases the initial magnitude and rate of depolarization.

**Figure 1.**
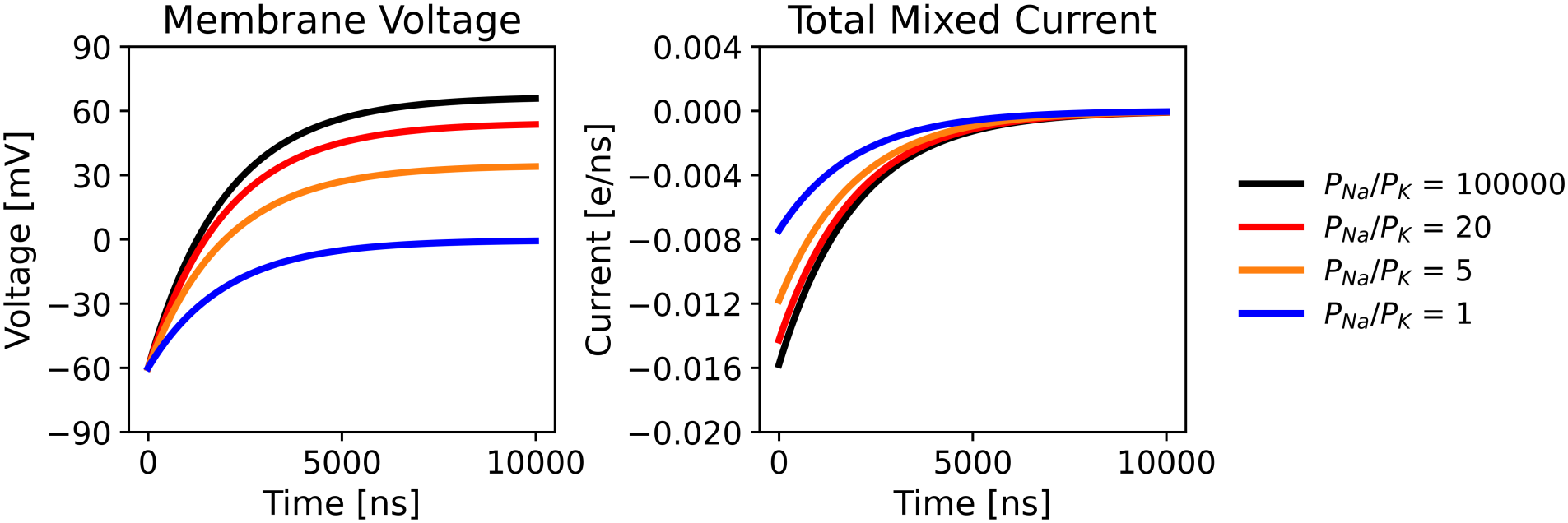
Results from the depolarization-only model. (A) Membrane voltage and (B) total mixed current as functions of time for P_Na_/P_K_ equal to 100000, 20, 5, and 1.

To understand the origin of this trend, Figure 1B shows the total mixed current through the conduction pathway. Because the fully Ohmic mixed current cannot be uniquely decomposed into ion-specific Na^+^ and K^+^ components, only the total mixed current is reported. We used a single conductance, G_mix_ = 20 pS, for the mixed-permeability pathway across all selectivity conditions, rather than separate Na^+^ and K^+^ conductances. With conductance held fixed, differences in current and membrane depolarization among conditions arose from selectivity-dependent changes in the GHK-calculated mixed reversal potential. Increasing P_Na_/P_K_ shifted the reversal potential toward the Na^+^ equilibrium potential and consequently increased the inward electrochemical driving force at the initial membrane voltage. This larger driving force increased the magnitude of the inward current, thereby charging the membrane more rapidly. As the membrane depolarized and approached the mixed reversal potential, the driving force decreased and the current gradually approached zero.

### Effect of Na^+^ channel selectivity in the modified Hodgkin-Huxley model

We next examined whether the same dependence of membrane depolarization on selectivity is preserved in the modified Hodgkin-Huxley model when the Na^+^ channel is assigned a finite Na^+^-to-K^+^ permeability ratio. Figure 2A shows the membrane voltage traces for four Na^+^ channel permeability ratios. For each selectivity, the initial membrane voltage was obtained by numerically solving the zero-current, steady-state form of Eq. 6, with the gating variables set to their voltage-dependent steady-state values (see Numerical implementation). The resulting equilibrium membrane voltages were −65.00, −65.09, −65.18, and −65.41 mV for P_Na_/P_K_ = 100000, 12.8, 5, and 1, respectively, and were used to initialize the simulations. All four equilibria were stable. As the Na^+^ channel selectivity was reduced, the depolarizing upstroke became slower and the peak membrane voltage decreased. In contrast, at high Na^+^ selectivity, the model produced a rapid upstroke and a larger peak voltage. Thus, the same qualitative trend observed in the depolarization-only model is preserved in the action-potential framework.

**Figure 2.**
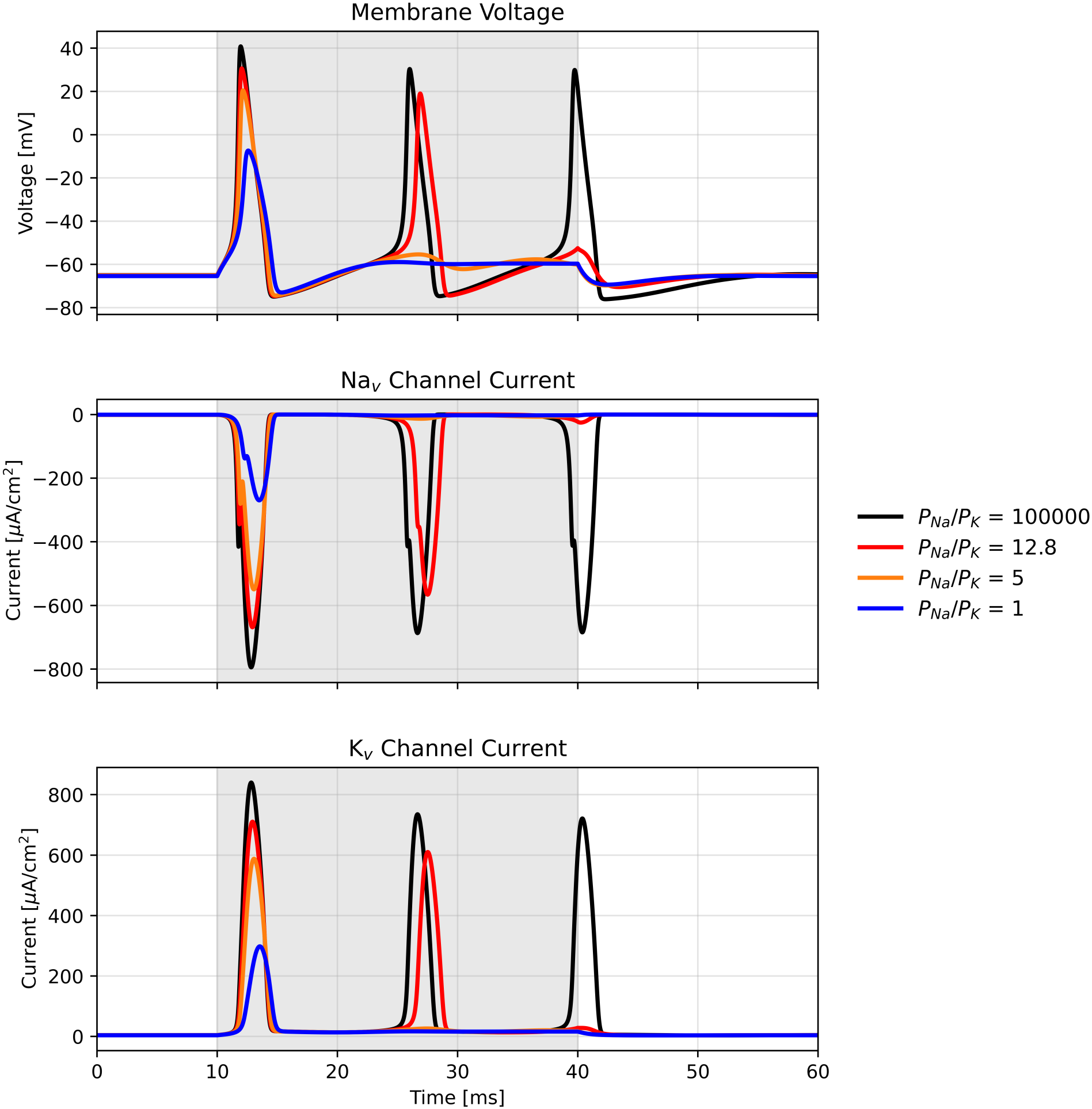
Effect of Na^+^ channel selectivity in the modified Hodgkin-Huxley model. (A) Membrane voltage, (B) total Na^+^ channel current, and (C) total K^+^ channel current for the indicated Na^+^ channel selectivity P_Na_/P_K_ equal to 100000, 12.8, 5 and 1.

The channel-current traces in Figure 2B/C clarify the mechanism underlying this behavior. Because the Na_v_ channels have finite Na^+^/K^+^ selectivity, the current shown in Figure 2B represents the total current through these channels, including contributions from both Na^+^ and K^+^ permeation. Figure 2B shows that reducing Na^+^ channel selectivity progressively decreased the magnitude of the inward total Na_v_ channel current. Because the maximal Na_v_ conductance and gating kinetics were unchanged, this reduction arose from the mixed GHK reversal potential.

Decreasing P_Na_/P_K_ shifted the Na_v_ reversal potential toward the K^+^ equilibrium potential, thereby reducing the inward driving force during depolarization. The smaller inward Na_v_ current produced a slower upstroke and a lower peak voltage. Figure 2C shows a corresponding reduction in the outward K_v_ channel current. Because the K_v_ channel formulation was identical across these simulations, this reduction was a secondary consequence of the weaker depolarization, which reduced K_v_ activation, as reflected in the lower n and n^4^ peaks (Figure S3A).

Na_v_ selectivity also affected repeated action-potential generation during the sustained current injection (Figure 2). At the highest selectivity, each large inward Na_v_ current transient initiated an action potential and was followed by a large outward K_v_ current that repolarized the membrane. As selectivity decreased, both the inward Na_v_ current transients and the resulting voltage responses became smaller, and the model generated progressively fewer full-amplitude action potentials. At the lowest selectivity (1), the inward Na_v_ current was insufficient to drive the membrane to positive voltages. Thus, lower Na_v_ selectivity weakens the depolarizing feedback required for both the action-potential upstroke and repeated action-potential generation.

To characterize the dependence of action-potential amplitude on Na_v_ channel selectivity, we performed a dense sweep of P_Na_/P_K_ from 0.3 to 100000. For each selectivity, peak membrane voltage was defined as the maximum voltage reached during the 10 to 40 ms current-injection interval. Figure 3 shows that peak voltage increased rapidly with P_Na_/P_K_ at low and intermediate selectivities and approached a plateau of approximately 41 mV at high selectivities. The experimentally reported squid-axon values of 3.5, 5.7, and 12.8 fall within the steep portion of the curve, indicating that changes in selectivity within this range can substantially affect action-potential amplitude, whereas further increases at high selectivity have relatively little effect.

**Figure 3.**
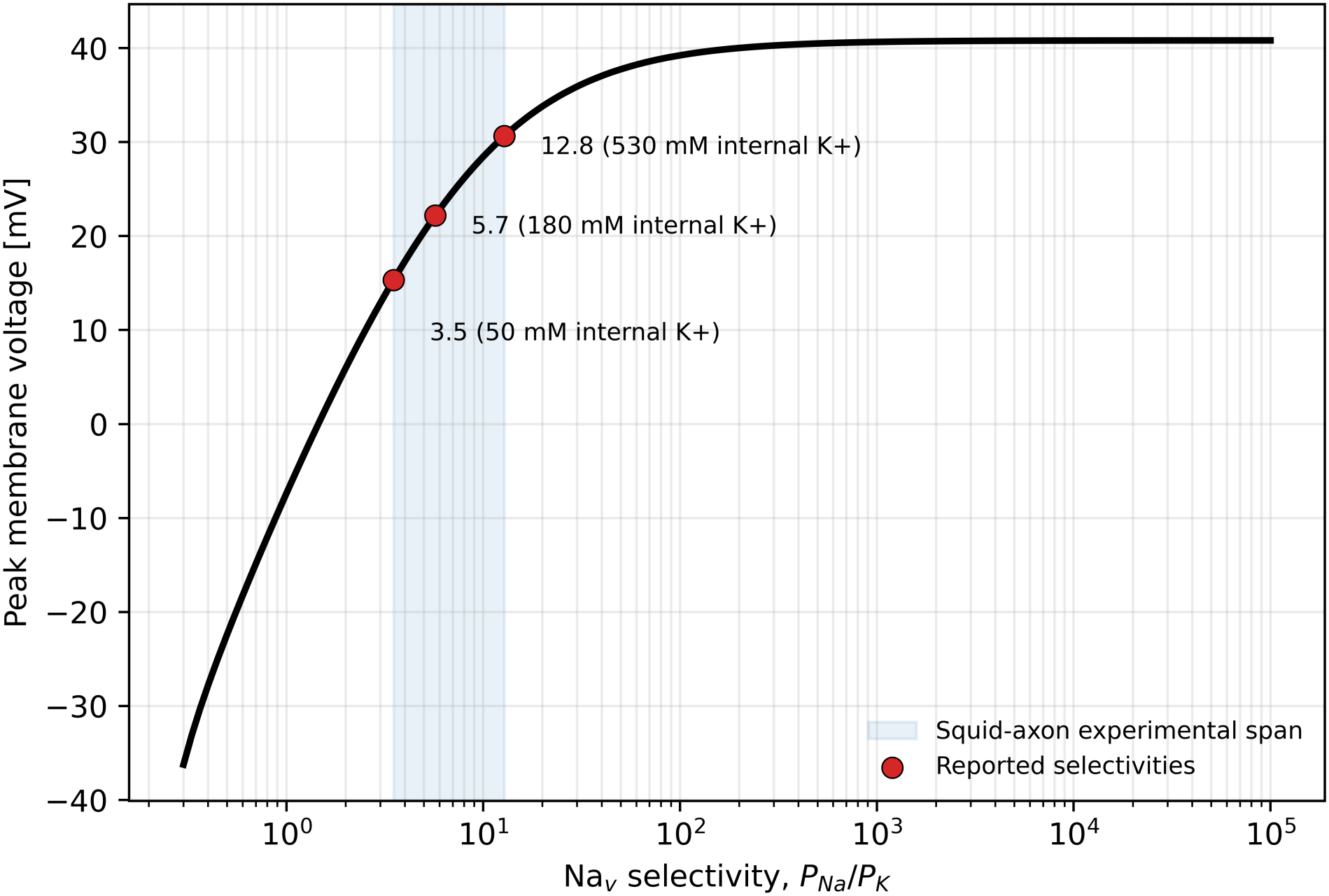
Dependence of action-potential peak voltage on Nav channel selectivity. The black curve shows the calculated peak voltage as a function of P_Na_/P_K_. The blue shaded region indicates the experimentally reported squid-axon selectivity range, and the red circles mark experimentally measured P_Na_/P_K_ values of 3.5, 5.7, and 12.8 obtained at intracellular K^+^ concentrations of 50, 180, and 530 mM, respectively.

### Effect of K^+^ channel selectivity in the modified Hodgkin-Huxley model

We then varied the selectivity of the K_v_ channel in the same modified Hodgkin-Huxley framework. Figure 4A shows the membrane voltage traces for different K_v_ channel permeability ratios. The effect of changing K_v_ channel selectivity was qualitatively different from that observed for the Na_v_ channel. Whereas Na_v_ channel selectivity mainly controlled the strength of the rapid depolarizing upstroke, reducing K_v_ channel selectivity shifted the mixed K_v_ reversal potential toward more positive voltages, reducing the outward driving force at a given membrane voltage and shifting the membrane equilibrium toward more depolarized voltages. Each simulation was initialized at its pre-calculated equilibrium membrane voltage obtained using the same steady-state zero-current calculation (Eq. 6): −64.99, −55.80, −28.41, and +0.03 mV for P_Na_/P_K_ = 10^-5^, 0.03, 0.2, and 1, respectively. At P_Na_/P_K_ = 10^-5^, the membrane generated repeated action potentials during stimulation and returned toward its stable equilibrium after stimulus withdrawal. At P_Na_/P_K_ = 0.03, spiking continued after stimulus withdrawal, consistent with the instability of its zero-current equilibrium. At P_Na_/P_K_ = 0.2 and 1, the membrane remained near its stable, depolarized equilibrium throughout the simulation and did not generate action potentials. Spontaneous firing at P_Na_/P_K_ = 0.03 was also observed when simulations started from a common −65 mV and included a 100 ms pre-stimulus equilibration interval (Figure S4).

**Figure 4.**
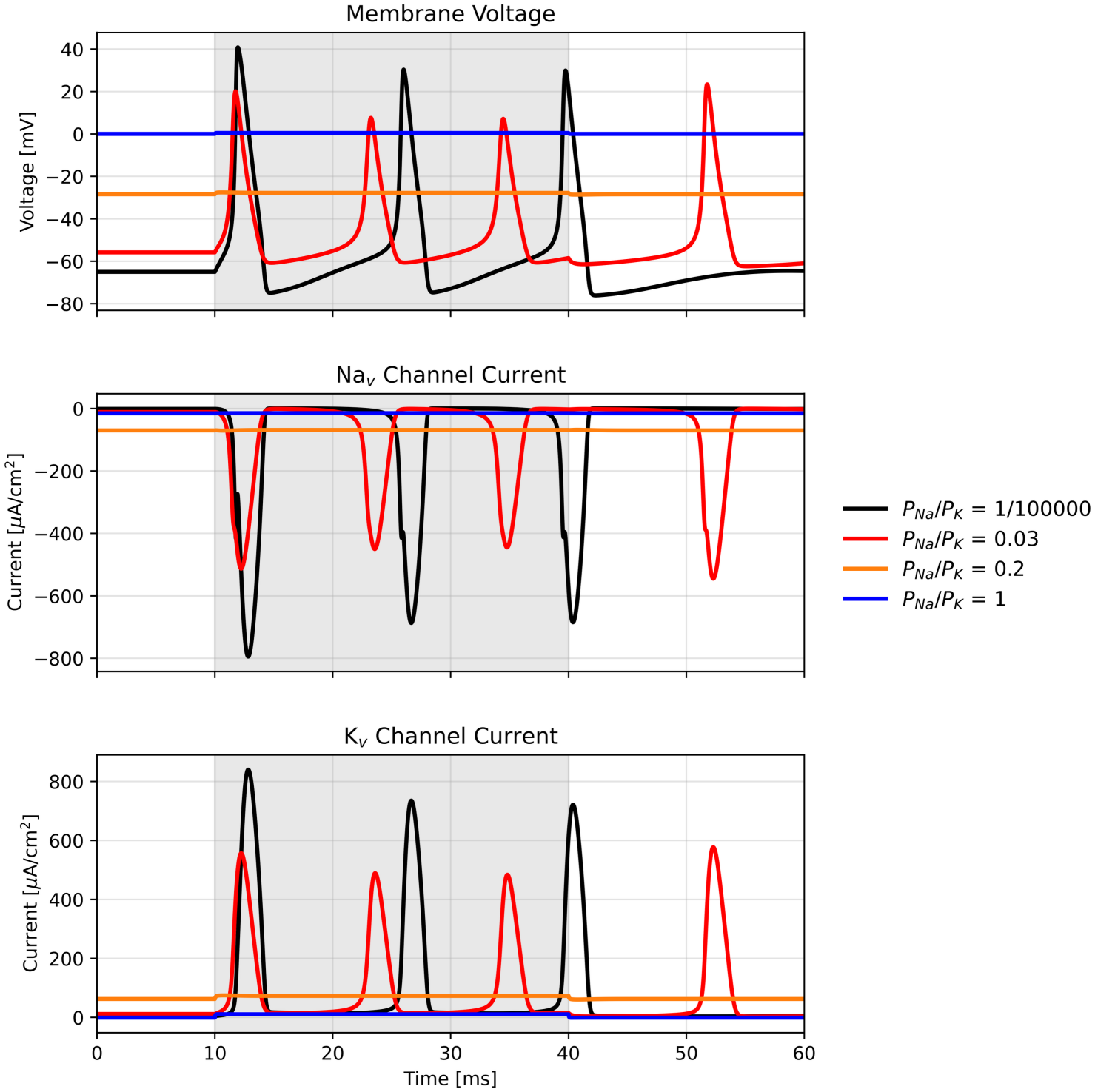
Effect of K_v_ channel selectivity in the modified Hodgkin-Huxley model. (A) Membrane voltage, (B) total Na_v_ channel current, and (C) total K_v_ channel current for the K_v_ channel selectivity P_Na_/P_K_ equal to 1/100000, 0.03, 0.2, and 1.

The total channel currents in Figure 4B/C and the gating variables in Figure S3B explain these different responses. At high K_v_ selectivity, each inward Na_v_ current transient was followed by a large outward K_v_ current that repolarized the membrane and allowed Na_v_ channels to recover from inactivation before the next action potential. At P_Na_/P_K_ = 0.03, the reduced outward K_v_ driving force weakened repolarization, leaving the membrane at less negative voltages between spikes. This limited Na_v_ recovery from inactivation, as indicated by the lower h values between action potentials and smaller m^3^h peaks (Figure S3B), contributing to the smaller inward Na_v_ current transients and action-potential peaks. At P_Na_/P_K_ = 0.2 and 1, the initial equilibrium voltages were already sufficiently depolarized to keep Na_v_ channels strongly inactivated. Despite high values of the activation variable m, the near-zero h values kept m^3^h small throughout the simulation (Figure S3B), preventing the large inward current transients needed for action-potential generation. Because the Na_v_ channel parameters were unchanged, these changes in Na_v_ current arose indirectly from the altered membrane voltage and its effects on channel gating.

To examine how K_v_ channel selectivity affects the membrane voltage after stimulation, we performed simulations over P_Na_/P_K_ values from 10^−5^ to 1 using the standard 12 µA/cm^2^ current pulse from 10 to 40 ms. Each simulation was run for 250 ms, and the mean and range of the membrane voltage during the final 50 ms were calculated. Figure 5 shows that the mean post-stimulus voltage remained close to −65 mV in the highly K^+^-selective regime but became progressively more depolarized as Na^+^ permeability increased. The widening voltage range at intermediate selectivities indicates that voltage oscillations persisted after stimulation rather than settling to a stable value, and the squid-axon upper-limit estimates of 0.03 and 0.06 fall within this strongly affected region. This overlap does not imply that normal squid axons would exhibit such oscillations, because finite Na^+^ permeability was introduced without recalibrating the other HH parameters originally fitted assuming K_v_ P_Na_/P_K_ = 0.

**Figure 5.**
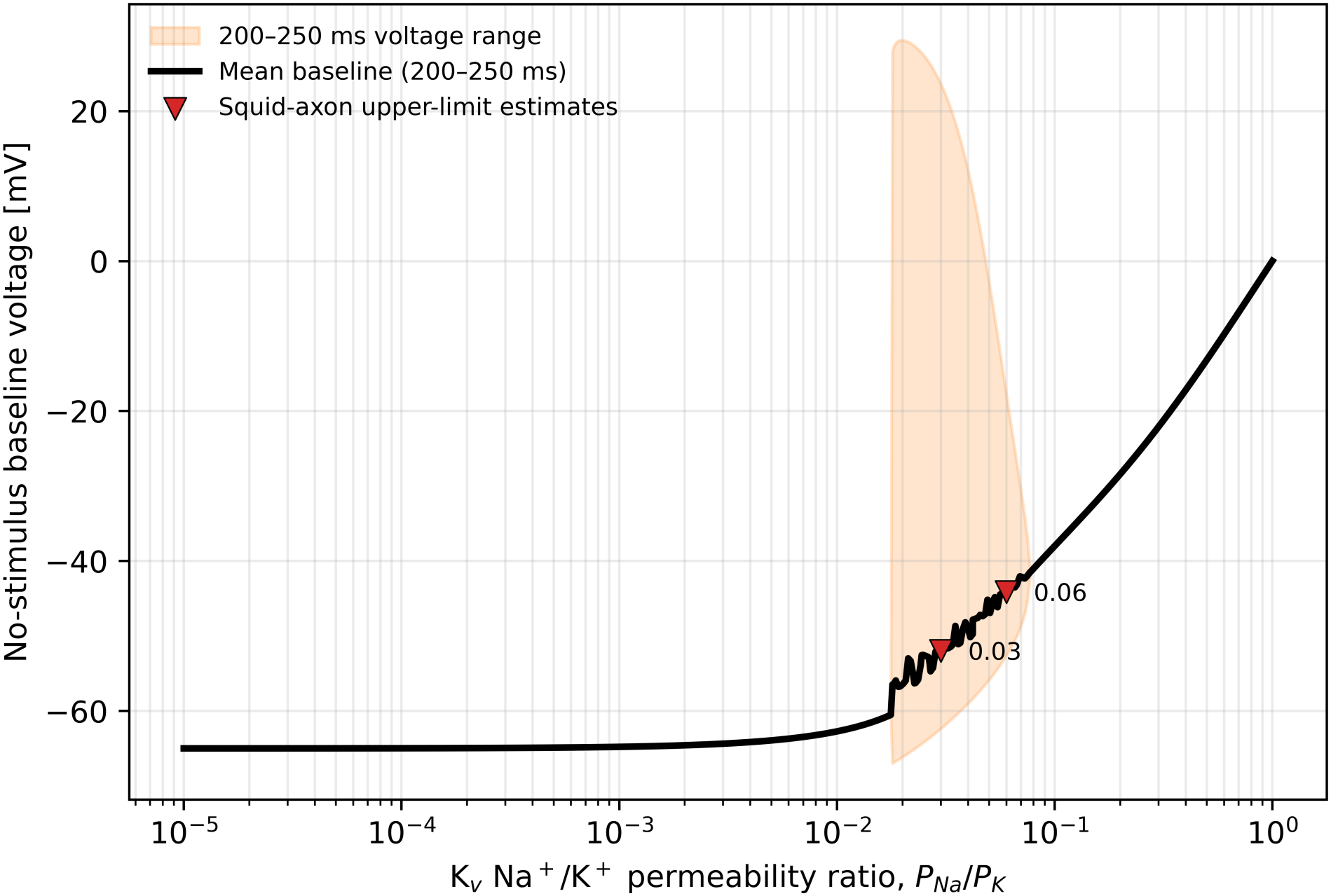
Dependence of post-stimulus membrane voltage on K_v_ channel selectivity. Simulations were run for 250 ms with a 12 µA/cm^2^ current pulse applied from 10 to 40 ms. The black curve shows the mean membrane voltage during the final 50 ms, and the orange shaded region shows the corresponding minimum-to-maximum voltage range. Red triangles indicate the squid-axon upper-limit estimates of P_Na_/P_K_ = 0.03 and 0.06.

### Effect of Na^+^ channel conductance in the modified Hodgkin-Huxley model

To separate the effect of selectivity from that of current magnitude, we next varied the maximal Na_v_ channel conductance while keeping the channel effectively fully Na^+^-selective. Figure 6 shows membrane voltage traces for maximal conductances of 60, 120, and 180 mS/cm^2^, with 120 mS/cm^2^ serving as the baseline used in the other simulations. Using the same steady-state zero-current calculation (Eq. 6), we obtained equilibrium membrane voltages of −65.47, −65.00, and −64.41 mV for maximal Na_v_ conductances of 60, 120, and 180 mS/cm^2^, respectively. These voltages were used to initialize the simulations, and all three equilibria were stable. At 60 mS/cm^2^, the model generated a smaller first action potential and did not produce additional action potentials during the sustained current injection. Increasing the conductance to 120 or 180 mS/cm^2^ produced larger, full-amplitude action potentials and supported repeated action-potential generation. The corresponding gating traces (Figure S3C) show repeated m^3^h peaks at 120 and 180 mS/cm^2^, whereas only the first large peak was present at 60 mS/cm^2^. Increasing conductance increased the magnitude of the inward current and accelerated membrane depolarization, resulting in earlier action-potential initiation. The effect on peak voltage was largely saturated between 120 and 180 mS/cm^2^, whereas the timing of action potentials remained sensitive to conductance. Thus, in addition to selectivity, the total available Na_v_ channel conductance strongly influences the initiation and repetition of action potentials.

**Figure 6.**
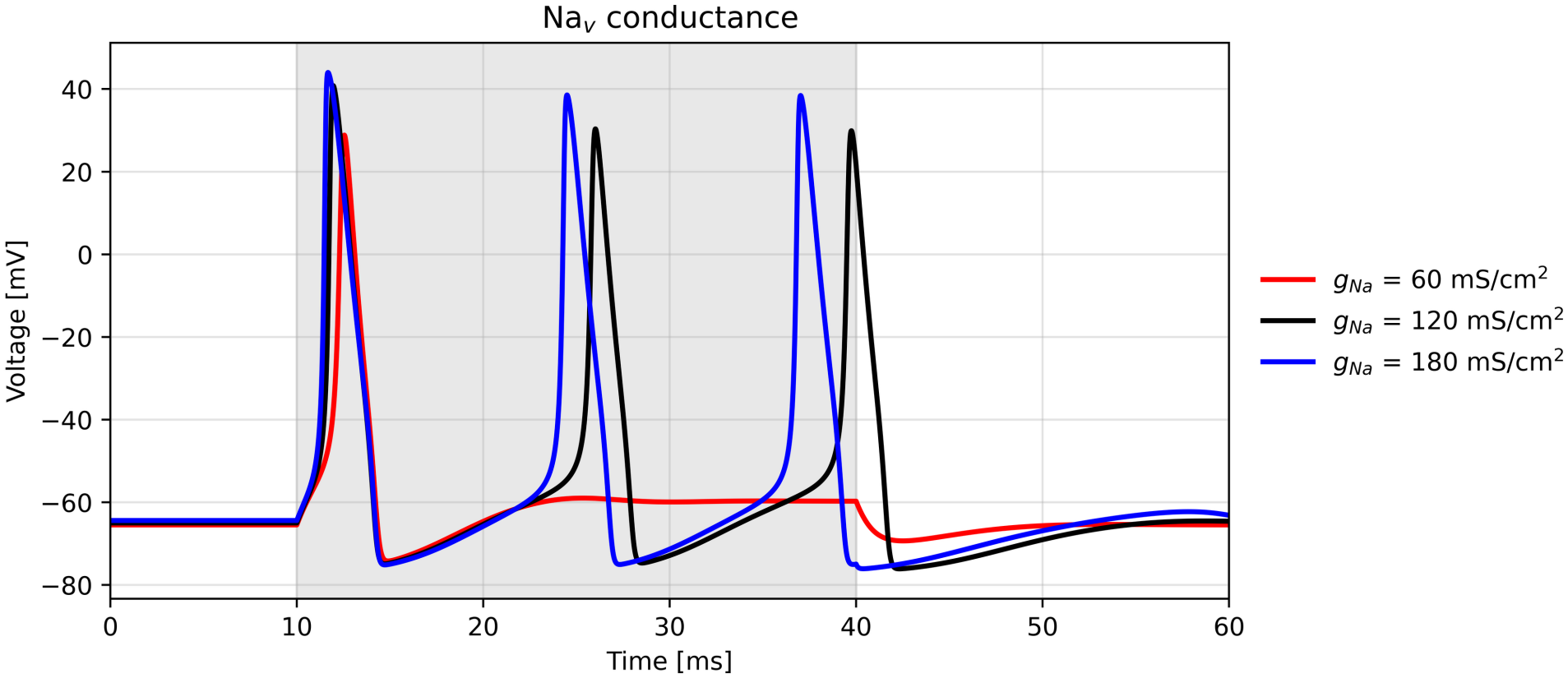
Effect of Na_v_ channel conductance in the classical Hodgkin-Huxley model. Membrane voltage traces for the maximal Na_v_ channel conductance equal to 60, 120 and 180 mS/cm^2^.

Comparison of Figure 6 with Figure 1 and Figure 2 shows that increasing Na_v_ conductance and increasing Na_v_ selectivity have qualitatively similar effects because both strengthen depolarization. However, they act through different parameters of the Ohmic current. Increasing conductance directly scales the total inward Na_v_ channel current without changing its reversal potential, whereas increasing selectivity shifts the mixed GHK reversal potential toward the Na^+^ equilibrium potential and thereby increases the inward driving force. Consequently, in the parameter ranges examined here, increasing conductance causes action potentials to occur earlier and promotes their repeated generation, while increasing selectivity primarily strengthens the depolarizing upstroke and increases peak voltage.

Taken together, the results of Figures 1 to 4 establish a consistent picture across the two models. Stronger Na_v_ channel selectivity shifts the mixed reversal potential toward the Na^+^ equilibrium potential, increases the inward driving force and for a given channel conductance and gating state thus increases the total inward current, and produces faster and stronger depolarization.

Reduced Na_v_ selectivity has the opposite effect and weakens the inward current driving the rising phase. In the modified Hodgkin-Huxley model, Na_v_ channel selectivity, K_v_ channel selectivity, and Na_v_ channel conductance affect membrane-voltage dynamics through distinct mechanisms: Na_v_ selectivity primarily controls the strength of depolarization, K_v_ selectivity determines the membrane voltage between or after action potentials and the ability to repolarize, and Na_v_ conductance influences the initiation and repetition of action potentials. Together, these properties influence the timing, rate of rise, amplitude, post-stimulus membrane voltage, and repetition of action potentials.

## Discussion

In this study, we examined how Na^+^/K^+^ selectivity influences action-potential dynamics using two complementary models: a simplified depolarization model and a modified Hodgkin-Huxley model. Both models showed that reduced ion selectivity alters membrane-voltage dynamics in a consistent direction. In the Hodgkin-Huxley model, reduced Na_v_ selectivity produced slower and smaller action-potential upstrokes and reduced repetitive activity, whereas reduced K_v_ selectivity impaired repolarization, shifted the membrane toward more depolarized potentials, and ultimately suppressed repetitive action-potential generation. Changes in Na_v_ conductance also altered action-potential dynamics, but through a mechanism distinct from changes in ion selectivity. These results show that ion selectivity and conductance represent separate biophysical properties that can independently shape membrane electrical behavior.

The different effects of Na_v_ and K_v_ selectivity arise from the different roles of these channels during the action potential. Under the ionic conditions used here, perfectly Na^+^-selective and K^+^-selective channels have reversal potentials of approximately +50 and −77 mV, respectively. For Na_v_ channels, decreasing P_Na_/P_K_ increases the contribution of K^+^ to the mixed reversal potential, shifting it from +50 mV toward more negative values, reducing the magnitude of V − E_rev_ during depolarization, and therefore decreasing the inward current that drives the action-potential upstroke. The resulting weaker depolarization reduces voltage dependent activation of K_v_ channels, as indicated by the smaller n^4^ peaks (Figure S3A), and also reduces the outward K_v_ driving force, so the decrease in K_v_ current (Figure 2) observed at lower Na_v_ selectivity is a secondary consequence of the altered voltage trajectory (Figure S3A). For K_v_ channels, reduced K^+^ selectivity corresponds to a decrease in P_K_/P_Na_, equivalently an increase in P_Na_/P_K_. This shifts the mixed reversal potential toward more positive values, reducing V − E_rev_ and therefore the outward current that drives repolarization. At sufficiently low K_v_ selectivity, the membrane remains depolarized between action potentials, keeping the Na_v_ inactivation variable h low and reducing the number of Na_v_ channels available for subsequent activation (Figure S3B).

The effects of varying Hodgkin-Huxley model parameters have been studied extensively, including changes in channel density or conductance, gating kinetics, ionic reversal potentials, and other properties that influence action-potential propagation, excitability, information transmission, and energetic cost (51, 52). Such studies have also asked whether experimentally observed parameter values occupy functionally favorable regions of the Hodgkin-Huxley parameter space. In particular, changes in reversal potentials E_Na_ and E_K_ have been shown to alter action-potential waveform, firing behavior, propagation velocity, and energetic cost (52). However, these studies did not treat Na^+^/K^+^ selectivity itself as an independent channel parameter. Reversal potentials were instead varied directly or through changes in ionic concentrations. Here, we calculate the mixed reversal potential from the relative Na^+^ and K^+^ permeabilities of the channel, allowing changes in ion selectivity to be propagated directly into membrane electrical dynamics. This formulation makes P_Na_/P_K_ itself the independent biophysical variable, while leaving the total channel conductance and gating formulation unchanged.

A similar question can be asked for ion selectivity: what range of relative permeabilities is sufficient to preserve characteristic action-potential dynamics? Experimentally measured selectivities vary among Na_v_ and K_v_ channels and can also be altered substantially by mutations within or near the selectivity filter. For example, classical measurements of Na_v_ channels in squid axons reported P_Na_/P_K_ values ranging from approximately 3.5 to 12.8 depending on intracellular K^+^ concentration (19), whereas measurements of K_v_ channels in squid axons reported upper limits on P_Na_/P_K_ of 0.03 and 0.06 (corresponding to lower limits on P_K_/P_Na_ of approximately 33 and 17, respectively) (48–50). These experimentally observed ranges provide a useful scale for determining when changes in selectivity become functionally important in the action-potential model. Our parameter sweep (Figure 3 and Figure 5) revealed that the dependence of action-potential properties on selectivity is strongly nonlinear. For Na_v_ channels, action-potential peak voltage increased rapidly with P_Na_/P_K_ over the range of experimentally measured selectivities and approached a plateau only at substantially higher P_Na_/P_K_ values. Thus, experimentally measured Na_v_ channels operate in a range in which relatively small fractional changes in selectivity can still influence action-potential amplitude. For K_v_ channels, increasing Na^+^ permeability progressively shifted the equilibrium membrane potential toward more positive values and impaired action-potential generation. More generally, these results show that the functional effect of a change in selectivity depends not only on the magnitude of the change but also on the initial selectivity of the channel.

At K_v_ P_Na_/P_K_ = 0.03, spontaneous firing persisted after stimulus withdrawal (Figure 4). Starting all conditions at −65 mV with a 100 ms pre-stimulus interval showed that the other three conditions approached their equilibrium voltages, whereas 0.03 produced repeated firing before and after current injection (Figure S4A), consistent with its computed zero-current fixed point being linearly unstable. Figure 5 shows that this behavior extends across an intermediate range of K_v_ selectivities. In this regime, Na_v_ inactivation and delayed K_v_ activation repolarize each spike sufficiently for h to recover and n to decline. The reduced outward K_v_ driving force then permits renewed depolarization and rapid m activation, sustaining the cycle without injected current (Figures S3B and S4B). This provides a selectivity-based counterpart to the conductance- and concentration-induced oscillations analyzed by Horng and Huang (2006) (53) and the reversal-potential-dependent stability transitions studied by Nguyen (2026) (54). Although 0.03 is experimentally motivated, the classical HH parameterization uses a K^+^-specific K_v_ current, equivalent to PNa/PK = 0. Without recalibrating the model to physiological squid axon behavior, introducing finite Na^+^ permeability while retaining the remaining parameters therefore perturbs that reference. Thus, the resulting instability does not itself invalidate either the experimental estimate or the classical model. Rather, it suggests a possible mechanism by which selectivity-altering mutations could promote abnormal firing, with neuron-specific parameters determining the relevant thresholds and physiological consequences.

An important feature of ion selectivity is that it can be related directly to the molecular properties of the ion channel pore. Relative Na^+^/K^+^ permeability and channel conductance can be measured experimentally and can also be calculated from molecular simulations (19, 26, 41, 55, 56). These quantities can be used to connect changes at the level of the channel pore to changes in ionic current and action-potential dynamics. In our previous work, we proposed a multiscale approach for linking molecular simulations to neuronal models (38), in which mutation-induced changes in channel conductance were propagated to changes in action-potential generation. The present study extends this connection to ion selectivity by showing how changes in Na^+^/K^+^ permeability ratio alter the reversal potential and consequently the ionic driving force, thereby affecting action potentials. Such an approach could be used to study the effects of molecular perturbations, including mutations, drug binding, and changes in protonation state or pH, on action-potential dynamics. Below, we illustrate this idea using the effect of pH on ion-channel conductance and selectivity.

Extracellular pH provides one example of a molecular perturbation that can influence ion-channel function. Changes in extracellular pH have been shown experimentally to affect the conductance of eukaryotic and bacterial Na_v_ channels, as well as voltage-gated Ca^2+^ channels (57–61). Such pH sensitivity may be relevant to neuronal signaling because extracellular pH undergoes small fluctuations (typically 0.1-0.2 pH units) associated with neuronal activity and metabolic state (62–67). In bacterial Na_v_ channels, acidification has additionally been shown to alter Na^+^/K^+^ selectivity (24), whereas experiments on eukaryotic Na_v_1.4 and Na_v_1.5 channels reported changes in conductance and gating without detectable changes in reversal potential (57). Our previous molecular simulations of a bacterial Na_v_ channel showed that protonation of selectivity-filter residues affects Na^+^ and K^+^ conductance differently, producing pH-dependent changes in both conductance and the Na^+^/K^+^conductance ratio (9, 55, 68), in agreement with experiments on bacterial channels. Using these molecular estimates, we explored how small changes in channel conductance and selectivity could influence action-potential dynamics (Figure S5). Because conductance and permeability ratios are not equivalent, and comparable pH-dependent changes in selectivity have not been established for eukaryotic Na_v_ channels, this analysis illustrates a possible molecular-to-cellular connection rather than providing a quantitative prediction of neuronal responses to pH.

In summary, our results show that Na^+^/K^+^ selectivity is an independent biophysical determinant of action-potential dynamics. Changes in Na_v_ and K_v_ selectivity alter their mixed reversal potentials and ionic driving forces, with distinct consequences for membrane depolarization and repolarization. Importantly, the effects of selectivity are nonlinear and differ between Na_v_ and K_v_ channels. For Na_v_ channels, further increases in P_Na_/P_K_ have relatively little effect once sufficiently high selectivity is reached, whereas channels within the experimentally observed selectivity range remain sensitive to changes in P_Na_/P_K_. For K_v_ channels, increasing Na^+^ permeability from the highly K^+^-selective regime progressively alters the equilibrium membrane potential and impairs repolarization. Because ion selectivity can be determined experimentally and calculated from molecular simulations, this relationship provides a direct route for propagating molecular changes in the channel pore to cellular electrical behavior. Incorporating ion selectivity into multiscale models may therefore help connect mutations, environmental perturbations, and other molecular modifications of ion channels to their functional consequences for electrical signaling.

## Acknowledgments

This work was supported by the Intramural Research Program of the NIH, NHLBI (ZIA HL001051-29). Ana Damjanovic acknowledges NHLBI Grant No. 75N92020P00042. Computational resources were provided by the LoBoS supercomputing cluster.

