## Supplementary Information for "Ion selectivity as a biophysical determinant of action-potential dynamics"

### Supplementary Methods

#### Depolarization only model with GHK currents

A GHK-current version of the depolarization-only model was used to quantify the intracellular concentration changes accompanying depolarization and assess the constant-concentration approximation in main text Figure 1. $Na^+$ and $K^+$ currents were calculated separately through the same mixed-permeability pathway using the GHK current equation (1, 2). For either monovalent cation i, the outward-positive current density was

$$J_i^{\mathrm{GHK}} = P_i \frac{F^2 V_m}{RT} \frac{[i]_{\mathrm{in}} - [i]_{\mathrm{out}} \exp\left(-\frac{F V_m}{RT}\right)}{1 - \exp\left(-\frac{F V_m}{RT}\right)}, \qquad i \in \{\mathrm{Na}, \mathrm{K}\} \qquad \text{Eq. S1}$$

Here, $P_{\mathrm{Na}}$ and $P_{\mathrm{K}}$ are absolute permeabilities; R, T, and F denote the gas constant, absolute temperature, and Faraday's constant. Permeability in m/s, voltage in V, and concentration in $mol/m^3$ (numerically equal to mM) give current density in $A/m^2$. Positive current denotes outward cation movement. The analytic limit was used at zero voltage.

For effective membrane area A, the ion-specific currents and transferred charges over one time step were:

$$I_{\mathrm{i}} = A J_{\mathrm{i}}^{\mathrm{GHK}}, \quad \Delta Q_{\mathrm{i}} = I_{\mathrm{i}} \Delta t \qquad \text{Eq. S2}$$

The membrane was represented as a capacitor, with capacitance C and voltage update:

$$C = \frac{\varepsilon_r \varepsilon_0 A}{L}, \quad V_{\mathrm{m}}(t + \Delta t) = V_{\mathrm{m}}(t) - \frac{\Delta Q_{\mathrm{Na}} + \Delta Q_{\mathrm{K}}}{C} \qquad \text{Eq. S3}$$

The relative dielectric constant $\varepsilon_r$ was 3.2, matching the main model and adopting the effective permittivity measured for DPPC bilayers in aqueous solution (3). Membrane thickness was 5 nm and the effective area per channel A was 0.007 $\mu m^2$, following the main model and its adopted channel-density scale (4). The negative sign follows the outward-positive current convention.

For a cylindrical axon, the ratio of intracellular volume to membrane surface area is R/2, where R is the axonal radius. Thus, we define the effective intracellular depth as $d_{\mathrm{in}}$ = R/2. Axonal diameters of approximately 1 to 1.5 µm have been reported for rat cortical layer 5 pyramidal neurons (4), corresponding to $d_{\mathrm{in}}$ = 0.25 - 0.375 µm. A representative value of $d_{\mathrm{in}}$ = 0.3 µm was chosen. Intracellular concentrations were updated in a well-mixed compartment of volume $Ad_{\mathrm{in}}$:

$$[i]_{\mathrm{in}}(t + \Delta t) = [i]_{\mathrm{in}}(t) - \frac{\Delta Q_{\mathrm{i}}}{FAd_{\mathrm{in}}}, \quad \mathrm{i} \in \{\mathrm{Na}, \mathrm{K}\} \qquad \text{Eq. S4}$$

Extracellular concentrations were fixed reservoirs. Voltage and intracellular concentrations were advanced by explicit forward Euler, with currents evaluated at the beginning of each step. The mixed reversal potential from main text Eq. 1 was a diagnostic only. The net GHK current vanishes at that voltage, although opposing ion-specific currents can remain nonzero.

The $Na^+$ permeability was calibrated from a 20 pS reference conductance in a symmetric 150 mM $Na^+$ solution at 310.15 K, matching the conductance scale adopted for the depolarization-only model in the main text:

$$P_{\mathrm{Na}} = \frac{g_{\mathrm{ref}} RT}{AF^2 C_{\mathrm{ref}}}, \quad P_{\mathrm{K}} = \frac{P_{\mathrm{Na}}}{S} \qquad \text{Eq. S5}$$

Using $g_{\mathrm{ref}}$ = 20 pS and $C_{\mathrm{ref}}$ = 150 mM gives $P_{\mathrm{Na}} = 5.276 \times 10^{-6}$ m/s. This calibration fixes $P_{\mathrm{Na}}$. The total GHK conductance is not constrained to remain constant as $P_{\mathrm{K}}$ changes. Simulations used similarity S = $P_{\mathrm{Na}}/P_{\mathrm{K}}$ = 100000, 20, 5, and 1.

Simulations started at -60 mV, with intracellular/extracellular $Na^+$ concentrations of 12/145 mM and intracellular/extracellular $K^+$ concentrations of 140/5 mM. The temperature was 310.15 K, the time step was 1 ns, and 10,000 steps were run, giving a total duration of 10 µs. No gating, leak, separate $K_v$ pathway, or injected current was included.

**Concentration variations in the Hodgkin Huxley model**

$Na^+$ and $K^+$ concentrations were varied one at a time in the classical Ohmic Hodgkin-Huxley model. Both channel reversal potentials were recalculated from the specified concentrations using the single-ion Nernst equation in main text Eq. 10 and used in the current equations (main text Eqs. 7 and 8). Concentrations were held constant within each run. Conductances, gating kinetics, leak parameters, and membrane capacitance were unchanged from the main Hodgkin-Huxley simulations.

The baseline concentrations were $[Na^+]_{in}$ = 55.17 mM, $[Na^+]_{out}$ = 440 mM, $[K^+]_{in}$ = 400 mM, and $[K^+]_{out}$ = 16.35 mM at 6.3 °C. The sweeps were $[Na^+]_{out}$ = 330, 440, and 550 mM; $[Na^+]_{in}$ = 40, 55.17, and 70 mM; $[K^+]_{out}$ = 12, 16.35, and 20 mM; and $[K^+]_{in}$ = 300, 400, and 500 mM. Each sweep included the baseline concentration and two integer concentrations approximately 25% below and above it. The other three concentrations retained their baseline values in each sweep.

Unless otherwise specified, Hodgkin-Huxley runs started at each condition's pre-calculated equilibrium membrane voltage, with gates initialized to $\alpha/(\alpha + \beta)$ at that voltage. The standard current pulse was 12 $\mu A/cm^2$ from 10 to 40 ms. Simulations used a 0.01 ms forward Euler time step and were run for 60 ms.

**pH dependent conductance ratios and action potential sensitivity**

The molecular conductance calculation for a bacterial $Na_v$ channel included three protonation states containing zero, one, or two additional protons, respectively, using the ion-dependent protonation model described in the main text (5). For each ion condition, the relative statistical weights were:

$$w_0 = 1, \; w_1 = 10^{pK_{a1} - \mathrm{pH}}, \; w_2 = 10^{pK_{a1} + pK_{a2} - 2\mathrm{pH}} \qquad \text{Eq. S6}$$

The protonation state fractions were obtained by dividing each weight by the sum of the three weights. The sequential $pK_a$ values were 6.9 and 6.7 in $Na^+$ solution and 7.5 and 6.7 in $K^+$ solution (5). Conductances of the zero-, one-, and two-proton states were 29, 12, and 0.5 pS for $Na^+$, and 39, 14, and 2.3 pS for $K^+$, respectively (5). The effective conductance in each ion condition was the state-fraction-weighted mean. The fractional change in $g_{\mathrm{Na}}/g_{\mathrm{K}}$ was calculated relative to pH 7.4.

The Hodgkin-Huxley sensitivity analysis independently decreased selectivity S = $P_{\mathrm{Na}}/P_{\mathrm{K}}$ by 4% or 7% from 3.5, 5.7, and 12.8. Each run started at its own pre-calculated equilibrium membrane voltage. Peak voltage was the maximum during the standard 10-40 ms current pulse. Changes were measured against the corresponding unperturbed reference. Only the magnitude of the fractional molecular conductance-ratio change motivated these perturbations, because conductance and permeability ratios are distinct quantities.

### Supplementary Figures

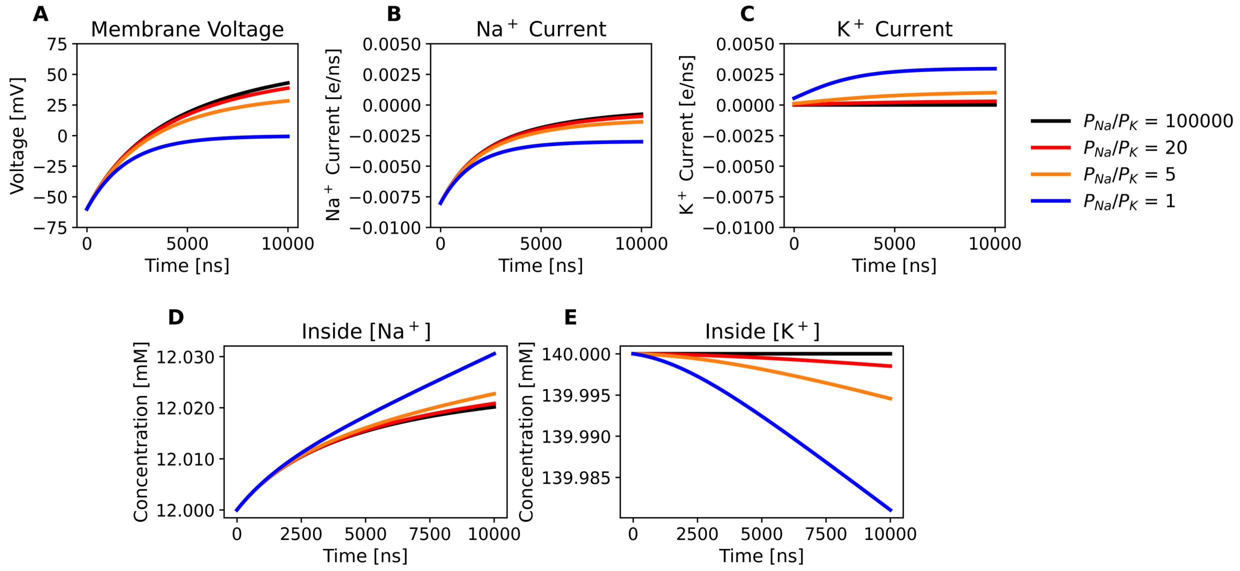


**Figure S1. Depolarization-only model with ion-specific GHK currents.** (A) Membrane voltage, (B) $Na^+$ current, (C) $K^+$ current, (D) intracellular $Na^+$ concentration, and (E) intracellular $K^+$ concentration for $P_{\mathrm{Na}}/P_{\mathrm{K}}$ = 100000 (black), 20 (red), 5 (orange), and 1 (blue). $Na^+$ permeability was fixed and $K^+$ permeability varied. Negative current denotes inward movement of positive charge; e/ns denotes elementary charges transferred per nanosecond. Extracellular concentrations were fixed. The maximum changes relative to the initial intracellular concentrations over the 10 μs interval were 0.255% for $Na^+$ and 0.0135% for $K^+$, supporting the constant-concentration approximation in main text Figure 1.

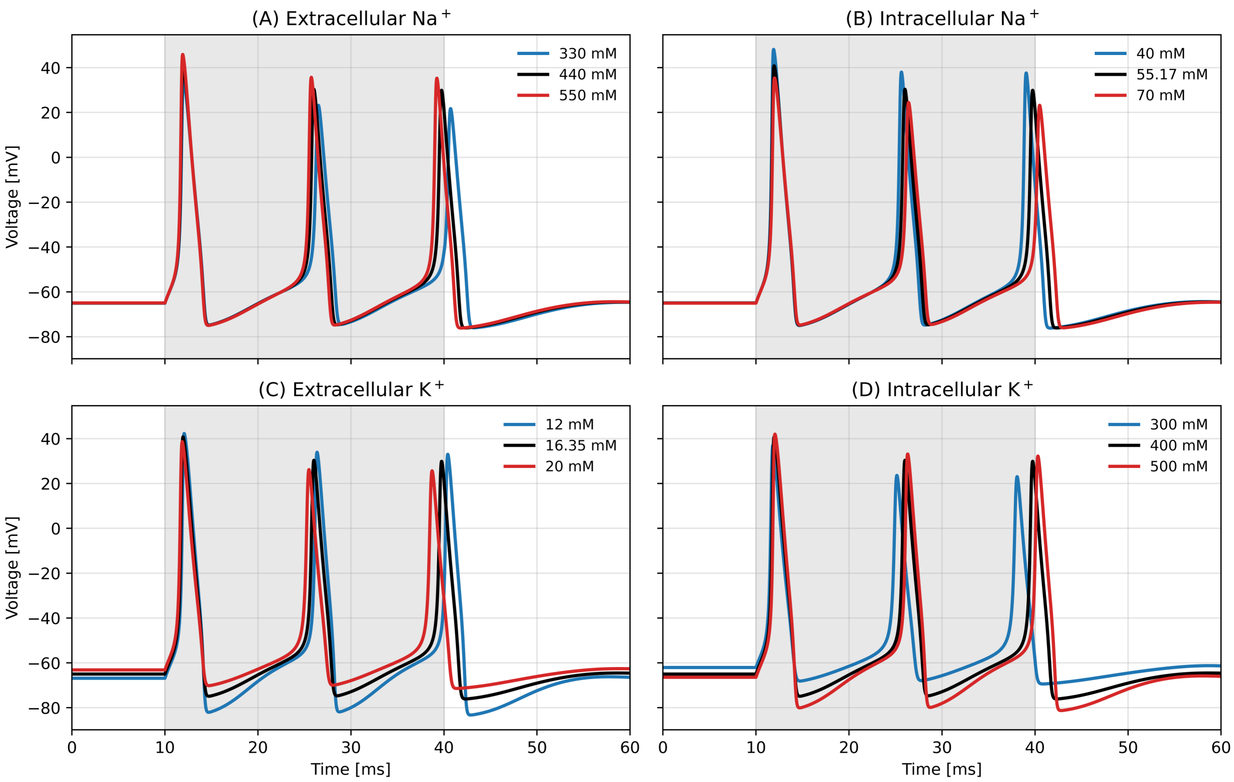


**Figure S2. Effects of ionic concentrations in the Hodgkin-Huxley model.** Membrane voltage traces for independent sweeps of (A) extracellular $Na^+$, (B) intracellular $Na^+$, (C) extracellular $K^+$, and (D) intracellular $K^+$. Concentrations are indicated in the legends; black curves represent the baseline condition. Only the indicated concentration changed in each sweep. Both channel reversal potentials were calculated with the Nernst equation. Each simulation began at its condition-specific equilibrium voltage. Gray shading marks the 12 µA/cm$^2$ current injection from 10 to 40 ms.

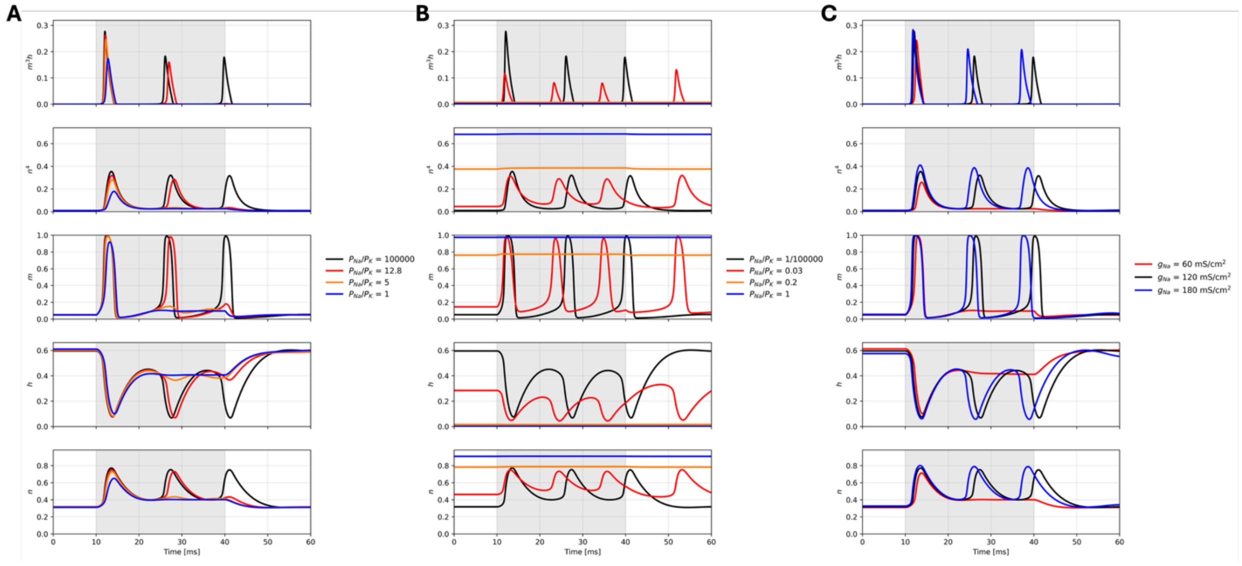

**Figure S3. Gating dynamics underlying the main Hodgkin-Huxley simulations.** Time courses for (A) $Na_v$ selectivity variations in main text Figure 2, (B) $K_v$ selectivity variations in main text Figure 4, and (C) $Na_v$ maximal-conductance variations in main text Figure 6. Rows show $m^3h$, $n^4$, m, h, and n, from top to bottom. The variables m and h describe $Na_v$ activation and availability, respectively, and n describes $K_v$ activation; $m^3h$ and $n^4$ are the corresponding conductance factors. Colors identify the conditions in each legend and match the corresponding main text figures. Gray shading marks current injection from 10 to 40 ms.

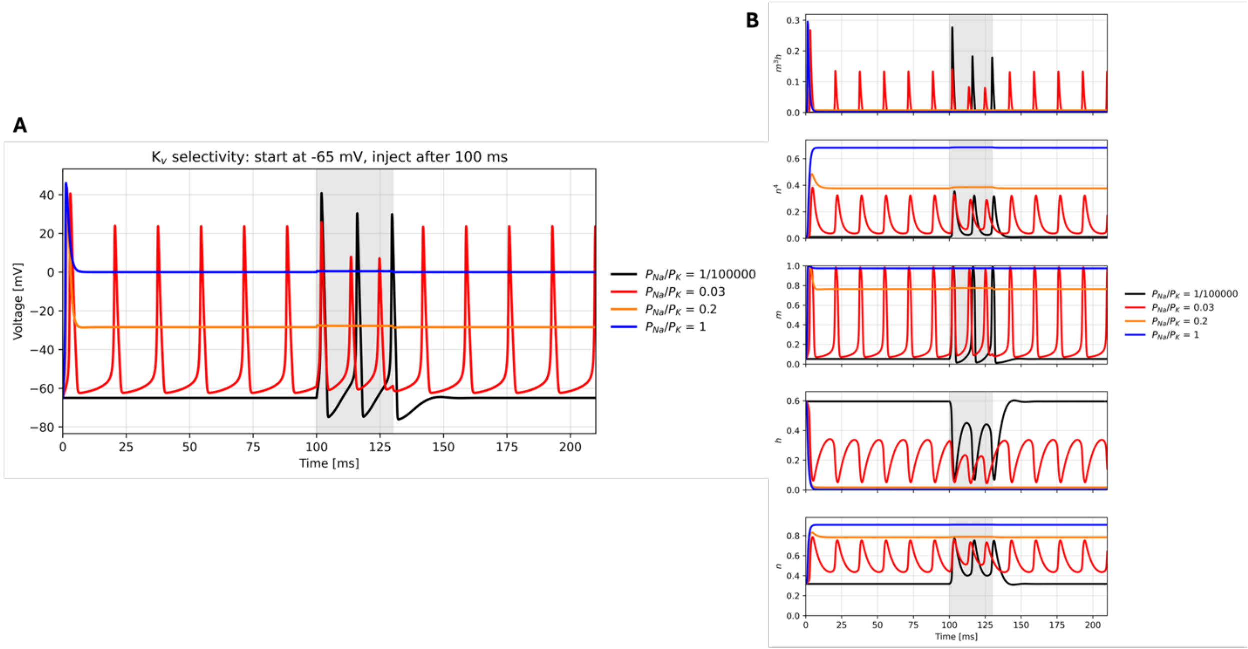


**Figure S4. $K_v$ selectivity with a common starting voltage and a 100 ms pre-stimulus interval.** (A) Membrane voltage and (B) the gating variables and conductance factors. All runs started at -65 mV with gates at their steady-state values at that voltage. A 12 μA/cm$^2$ pulse was applied from 100 to 130 ms (gray shading). $K_v$ $P_{\mathrm{Na}}/P_{\mathrm{K}}$ was 1/100000 (black), 0.03 (red), 0.2 (orange), or 1 (blue). At 0.03, reduced repolarizing drive makes the zero-current equilibrium voltage unstable. Repetitive firing persisted before and after injection: delayed $K_v$ activation repolarized each spike sufficiently for h to recover and $Na_v$ activation to recur. The 100 ms pre-stimulus equilibration interval therefore did not produce a stable resting voltage. At 0.2 and 1, sustained depolarization kept h near zero and suppressed subsequent full-amplitude action potentials.

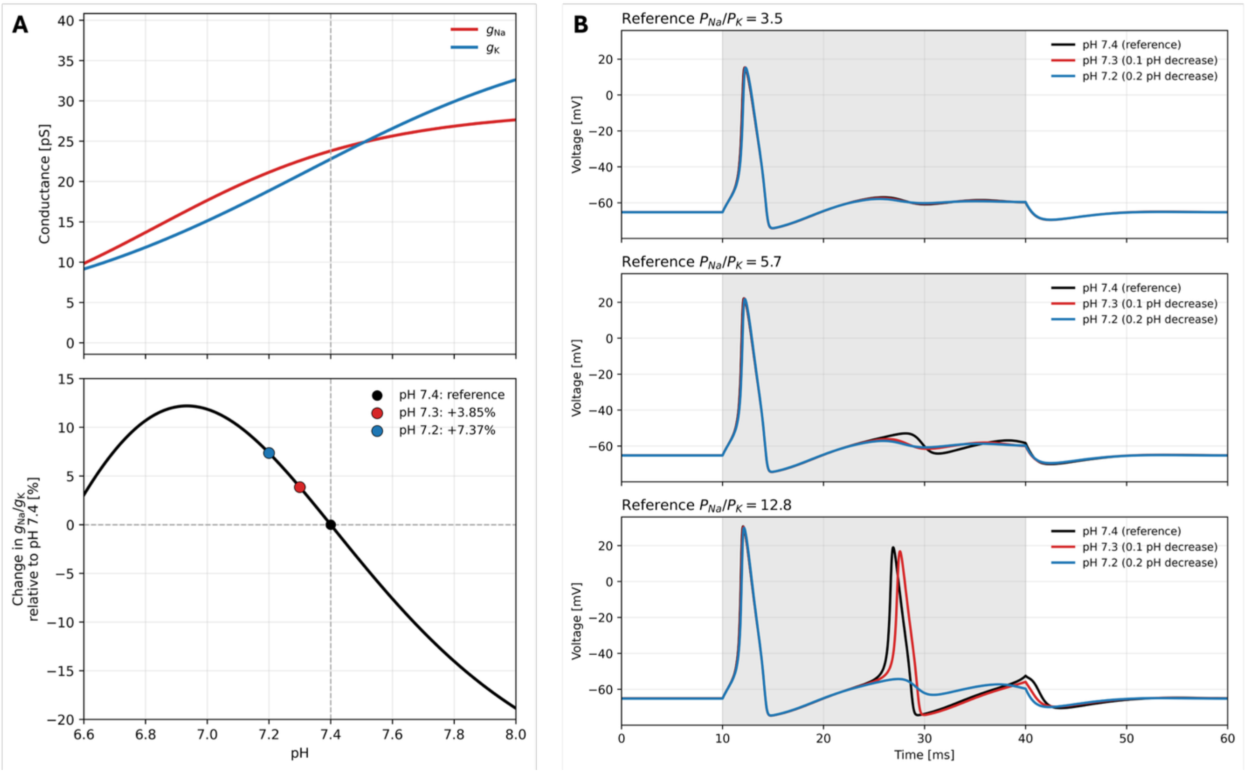


**Figure S5. pH-dependent channel conductance and selectivity and their effects on action potentials.** (A) Calculated Na$^+$ and K$^+$ conductances through a bacterial Na$_v$ channel (top) and percentage change in their ratio relative to pH 7.4 (bottom), using the protonation-state populations and state-specific conductances described in Supplementary Methods. Decreasing pH from 7.4 to 7.3 or 7.2 increased the conductance ratio by 3.85% or 7.37%, respectively. (B) Hodgkin-Huxley membrane voltage traces for baseline Na$_v$ $P_{\mathrm{Na}}/P_{\mathrm{K}}$ values of 3.5, 5.7, and 12.8 (top to bottom), at pH 7.4 (black), 7.3 (red), and 7.2 (blue). Assuming that eukaryotic Na$_v$ channels share the fractional pH responses of the bacterial channel, Na$_v$ maximal conductance and selectivity were changed simultaneously: conductance was scaled by the molecular Na$^+$ conductance change, while the conductance-ratio change in (A) was used as a proxy for the fractional permeability-ratio change. Peak membrane voltage during current injection changed by -0.723 to +0.059 mV relative to the corresponding pH 7.4 reference. Gray shading marks the 12 μA/cm$^2$ current pulse from 10 to 40 ms.
